# Polyandry and paternity affect disease resistance in eusocial wasps: support for the parasite–pathogen hypothesis

**DOI:** 10.1101/869537

**Authors:** Tatsuya Saga, Masaki Okuno, Kevin J. Loope, Koji Tsuchida, Kako Ohbayashi, Masakazu Shimada, Yasukazu Okada

## Abstract

Polyandry (multiple mating by females) is a central challenge for understanding the evolution of eusociality. Several hypotheses have been proposed to explain its observed benefits in eusocial Hymenoptera, and one, the parasite–pathogen hypothesis, proposes that high genotypic variance among workers for disease resistance prevents catastrophic colony collapse. We tested the parasite–pathogen hypothesis in the polyandrous wasp *Vespula shidai*. We infected isolated workers with the entomopathogenic fungus *Beauveria bassiana* and quantified their survival in the laboratory. Additionally, we conducted paternity analysis of the workers using nine microsatellite loci to investigate the relationship between survival and matriline/patriline of the workers. As predicted by the parasite–pathogen hypothesis, nestmate workers of different patrilines showed differential resistance to *B. bassiana*. We also demonstrated variations in virulence among strains of *B. bassiana*. Our results are the first to directly support the parasite–pathogen hypothesis in eusocial wasps and suggest that similar evolutionary pressures drove the convergent origin and maintenance of polyandry in ants, bees, and wasps.

## Introduction

Explaining the adaptive significance of polyandry (multiple mating by females) in eusocial insects has been a central challenge in sociobiology for the past three decades (Crozier and Pamilo 1996). However, since polyandry causes dilution of kin relationships among nestmates, the phenomenon seems contradictory to kin selection theory because high intra-colony relatedness would appear necessary for the evolution of eusociality (Crozier and Pamilo 1996; Queller and Strassmann 1998). Despite there being cost factors to female multiple mating (e.g., time and energy costs, Thornhill and Alcock 1983; predation risk, Arnqvist 1989; pathogen or virus transmission risk, Sherman et al. 1988; Amiri et al. 2016), polyandry and polygyny (presence of multiple queens in a nest) have evolved from ancestral monandry independently in eusocial insects, suggesting selective forces favoring genetic diversity among nestmates (Hughes et al. 2008). Large-scale phylogenetic studies have demonstrated that increased colony size is associated with greater paternity frequency and reduced paternity skew, both of which increase intra-colony genetic diversity across species of eusocial Hymenoptera (Jaffé et al. 2012; Loope et al. 2014).

Several plausible hypotheses have been proposed to explain the evolution of multiple mating given these costs (e.g., parasite–pathogen hypothesis, facilitating division of labor hypothesis, and gaining sperm hypothesis, reviewed in Crozier and Fjerdingstad 2001). The parasite-pathogen hypothesis (PPH) invokes the benefits of high intra-colony genetic diversity to explain the evolution of polyandry in social insects (Sherman et al. 1988; Shykoff and Schmid-Hempel 1991). The PPH assumes that different matrilines or patrilines of workers within the colony have different susceptibility or resistance to a single pathogen genotype, and also that different pathogen genotypes have different virulence to individuals of matrilines or patrilines (Sherman et al. 1988; Kraus and Page 1998; Sherman et al. 1998). The PPH proposes that the increased intra-colony genetic diversity arising from polyandry could prevent catastrophic colony collapse following challenge by parasites and pathogens, particularly if resistance to them among workers is variable depending on genotype (Sherman et al. 1988). The PPH has been supported only in Apidae and Formicidae, in which families polyandry has originated independently (Baer and Schmid-Hempel 2003; Palmer and Oldroyd 2003; Hughes and Boomsma 2004; Seeley and Tarpy 2007).

Here, we conducted two experiments to test the PPH in a eusocial wasp in Japan. The first aimed to evaluate fungal virulence differences between colonies, and the second aimed to evaluate fungal virulence differences among patrilines, worker emergence date, and their interaction in a more detailed fashion. We challenged multiple patrilines and matrilines of *Vespula shidai* with multiple strains of the entomopathogenic filamentous fungus *Beauveria bassiana. B. bassiana* infects a broad range of insect hosts, and one of its identifying features is the formation of its characteristic conidiospore balls (Hajek and Leger 1994; Schmid-Hempel 1998; Schmidt et al. 2011).

The nesting biology of *V. shidai* is similar to that of other *Vespula* (Matsuura 1995; Saga et al. 2017). A new colony is initiated by a single foundress queen that emerges from hibernation in spring. After hibernation, the queen founds the colony and produces workers and the colony grows through the summer, switching to the production of reproductives (males and new queens) from the beginning of fall to the beginning of winter. Mating occurs in the fall, and after the beginning of winter (early December in central Japan), only gynes hibernate, and all workers and males die (Matsuura & Yamane 1990).

In addition, we investigated whether differences in the growth environment affected resistance to disease, and whether workers that emerged on the same day showed any bias toward specific patrilines. Assuming that the benefits of high intra-colony genetic diversity are the evolutionary cause of polyandry, simultaneous emergence of workers from various patrilines would increase the genetic diversity within the nest. In many organisms, the effects of epigenetics and phenotypic plasticity mean that even individuals of the same genotype exhibit variability in disease resistance due to differences in environmental factors (Pigliucci 2005). In addition, given the homeostasis common within advanced eusocial societies (Oldroyd and Fewell 2007), it seems likely that workers emerging contemporaneously would have experienced very similar environments as larvae (e.g., kind of food and exposure to diseases).

The PPH has not yet been directly tested in *Vespula*, a genus phylogenetically independent of Apidae and Formicidae. In this study, our aims were to first test the prediction that matrilines/patrilines differ in their resistance to particular strains of pathogen, and then to determine if pathogen strains vary in their virulence to hosts of the same genotype, which would suggest the potential for classic co-evolutionary host–pathogen dynamics (Schmid-Hempel and Ebert 2003). We also hoped to extend the sparse literature available (Barribeau et al. 2014) on variation in pathogen strain virulence to the same eusocial wasp genotype host.

## Material and Methods

### Pathogens

We used five fungal strains (A–E) of the entomopathogenic filamentous fungus *Beauveria bassiana* for experiment 1, and then selected the two most lethal strains (A and C) for experiment 2. The fungal strains were isolated from five mummy queen larvae from different colonies of *V. shidai* collected in Nakatsugawa City, Gifu Prefecture, Japan, November 3, 2013. The strains were stored on Sabouraud agar medium (0.01 g/mL yeast extract, 0.01 g/mL bacto peptone, 0.02 g/mL L-glucose, 0.02 g/mL agar, and distilled water) in a refrigerator. The fungus was cultured on Sabouraud medium three times prior to the experiments. Conidiospore suspensions were prepared from sporulating culture plates (cultured at 25 °C, 8–14 days, 24-h light) in a 0.03% Tween 80 solution.

The fungal spore suspension (30 mL) was quantified using a hemocytometer (Sigma, St. Louis, MO, USA) and diluted to the required dosage for both experiments: low (10^6^ spores/mL) and high (10^8^ spores/mL).

### Wasps and Experiment Set-up

#### Experiment 1: Interaction of matriline and pathogen strain

We collected about 400 workers from each of five colonies, on November 9, 2014 at Akechi Wasp Festival in Toyota City, Aichi prefecture, Japan (regarding wasp festivals, refer to Nonaka 2010). On the day of collection, the workers were confined within wire mesh cages, one cage per colony, and placed into a room at constant temperature (25 °C with a 15-h light:9-h dark photoperiod). A 30% honey solution was provided daily for one week from the collection date, and the workers remaining alive were used in experiment 1. The workers were placed in a freezer at −20 °C for 20 s to anesthetize them just before conidiospore infection. Individual workers were then completely submerged for 1 s into either into the high dose fungal spore solution (10^8^ spores/mL, 30 mL) or into sterile water (30 mL; control). Excess fluid on the workers’ body surface was absorbed using filter paper. Each worker was then housed singly in a plastic box (3 × 8 × 10 cm) lined with filter paper, and provided with 500 μL water daily. Their survival was checked daily for 7 days, and their recorded survival time (in days) was recorded. If a worker died, we moisturized the filter paper in the box with water to maintain humidity at 90% for two more weeks. We attributed death to *B. bassiana* where whitish hyphae, a typical feature of the fungus, were seen growing from the corpse.

#### Experiment 2: Comparison of strain virulence between different worker patrilines and emergence dates

We collected three colonies in Nakatsugawa City, Gifu prefecture, Japan in middle of July 2015 using traditional methods for wasp hunting and keeping (Saga 2019), and labeled them TK, TG, and KM. We placed each colony in a wooden box (30 × 30 × 60 cm) and provided the colonies with meat (purchased chicken heart) and a 30% honey solution, permitting free outdoor foraging within the original collection areas until October 11, 2015. On this day, we collected combs filled with larvae and pupae from each colony, put the combs in a plastic box with a mesh roof, and set the boxes in a room at constant temperature (25 °C with a 15-h light:9-h dark photoperiod). Every 12 h, we transferred newly emerged workers to individual plastic boxes (3 × 8 ×10 cm) lined with filter paper. We conducted experiment 2 using the workers 12–24 h post emergence, administering the low concentration of fungal spore suspension (10^6^ spores/mL; 30 mL) of either strain A or strain C, using the same infection protocol as in experiment 1, including controls. Strains A and C were used because their apparent virulence differed most (Table 1). After inoculation, we placed the workers into individual boxes and fed them with 30% honey solution (2 mL) every 48 h, checking for death every 12 h for 15 days. Following death, we collected the worker’s antennae and middle legs for genotyping. As in the preliminary experiment, high humidity was maintained in the box to confirm that cause of death was *B. bassiana*.

**Table 1.**
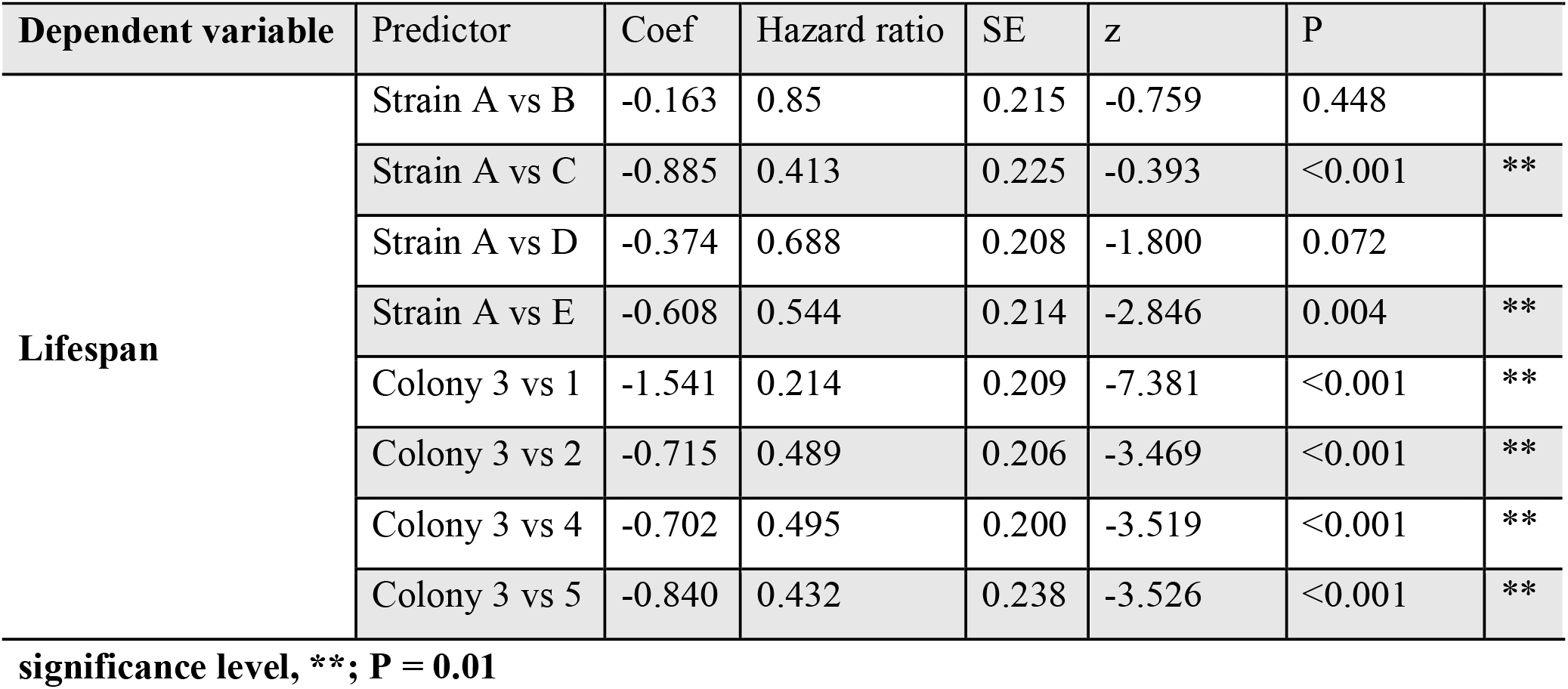
Cox survival analysis investigating the relationship between the lifespan and the strain used for infection, the colony, and the interaction of the both.

### Genetic analysis

We extracted DNA from the workers killed by *B. bassiana*. Template DNA was extracted from antenna or leg of the individuals by placing them in 50 μL of 5% Chelex solution (Chelex 100, 100–200 mesh, Bio Rad) and 0.5 μL of proteinase K (20mg/mL, Takara Bio Inc.), then incubating for 24 h at 56 °C and 5 min at 95 °C. We froze 5 μL of the supernatant in 45 μL of Tris-EDTA buffer (10 mM Tris, 1 mM EDTA, Takara Bio Inc.) before analysis by polymerase chain reaction (PCR). We genotyped workers at nine polymorphic microsatellite loci: List2001, List2003, List2004, List2019, List2020, Rufa5, Rufa19, VMA3 and VMA6 (Thorén et al. 1995; Daly et al. 2002; Hasegawa and Takahashi 2002) using multiplex PCR. We used dye-labeled primers (Applied Biosystems) in combination with a 3-primer labeling method (Schuelke 2000) to perform multiplex PCR with nine primers. Each 10 μL PCR reaction included 1 uL extracted DNA, 5 μL Qiagen master mix (Qiagen Type-It Microsatellite Kit, Qiagen Inc.), 0.2 μL of each reverse primer, 0.2 μL (dye-labeled) or 0.1 μL (3-primer labeled) of each forward primer, 0.15 μL FAM-labeled 3-primer tag for each 3-primer-labeled primer pair, and water to total 10 μL. PCR reaction conditions were 95 °C for 15 min, 35 cycles at 95 °C for 30 s, 50 °C for 90 s, 72 °C for 60 s, followed by 60 °C for 30 min. Fragment analysis was performed on an ABI-3730 × l sequencer using 0.5 μL PCR product combined with 15 μL HiDi Formamide and 0.15 μL LIZ 500 internal size standard (Applied Biosystems). Allele sizes were called using GeneMarker (SoftGenetics LLC) and checked twice by eye. We used 10 samples to check for linkage, departures from Hardy–Weinberg equilibrium, and null alleles using Genepop 4.4 (Rousset 2008; Supplementary Materials and Table S1).

### Estimating maternity and paternity

We used nine microsatellite markers which were suitable for estimating maternity and paternity (see Supplementary Material). To determine patriline membership, we analyzed microsatellite genotypes using the software Colony v2.0.6.5 (Wang 2004). First, genotypes at locus L2001 were set to unknown for individuals from colony TK because genotypes suggested a maternal null allele. We then ran Colony using a maternal sibship constraint for each colony *(V. shidai* is an obligatory monogynous species). We then reexamined genotypes that were identified as possibly erroneous by Colony, and also rescored genotypes for individuals whose putative fathers’ genotypes differed from other fathers at only a single locus, as these are likely the result of genotyping errors. Colony was then re-run using the corrected genotypes to produce the final sibship assignments. Settings for the Colony runs were: updating allele frequencies, inbreeding absent, polygamy for females, monogamy for males, no scaling of full sibship, no sibship size prior, unknown allele frequencies, a long full likelihood run, and 0.01 dropout and other error rates for all loci.

Sibship assignments using Colony identified 3–4 major patrilines per colony. One worker in colony TK and two in colony KM were not assigned to one of the common patrilines, and likely represented minor patrilines, foreign workers, or genotyping errors. These individuals were removed from further analyses.

### Statistical analysis

For experiment 1, we compared each of the Kaplan–Meier survival curves between the infection and control treatment by log-rank test. We also tested the influence of each colony and each strain, and interaction between both on lifespan checked every 24 h by Cox regression survival analysis with variable selection by stepwise method (stepAIC function, MASS package, R). For Cox regression survival analysis we used colony and strain, which had the lowest survival rate in experiment 1, as a reference for each comparison. All statistical analyses were conducted in R 3.2.4 (R 2018).

For experiment 2, we analyzed the relationship between worker emergence date and patrilines using ANOVA. We compared each of the Kaplan–Meier survival curves in a pair-wise fashion between the infection and control treatment by log-rank test. We also tested the influence of each patriline, each strain, each emergence date and their interactions on lifespan by Cox regression survival analysis with variable selection by stepwise method in each colony, since patrilines are nested within colonies (stepAIC function, MASS package, R). For Cox regression survival analysis we used the patriline that had the lowest survival rate in experiment 2 as a reference for comparison within each colony. All statistical analyses were conducted in R 3.2.4 (R 2018).

## Results

The survival time of controls was significantly longer than of the workers infected with each strain of *B. bassiana* (Table S2). The Akaike information criterion (AIC) value calculated for the full model of the results of experiment 1 (dependent variables: survival time and rate; predictor variable: each colony and each strain, the interaction of the both) was 1368.1. The AIC value calculated in the final model (dependent variables: survival time and rate; predictor variable: each colony and each strain) after performing stepwise model selection was 1351.3. Any interactions were excluded as a variable by the stepwise model selection. We found that differences of colonies and strains had a significant influence on lifespan in experiment 1 (Table 1). The results of experiment 1 are illustrated in Fig. 1.

**Figure 1.**
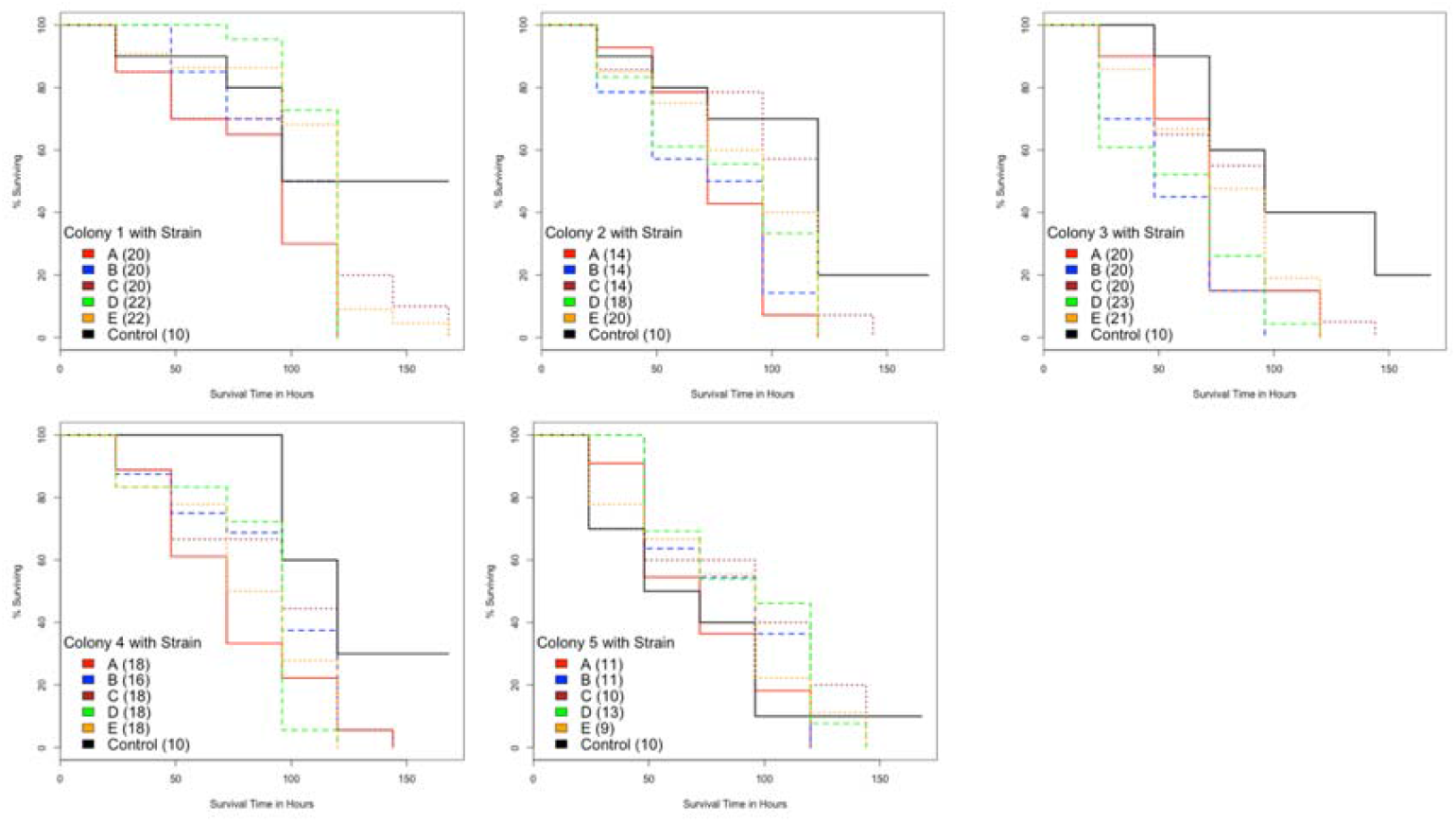
Survival curves for controls and infected workers in each colony. Legends show strains of *B. bassiana*. Values in parentheses are number of samples. Table 1 shows the statistical analysis of these results.

We did not detect any deviation from Hardy–Weinberg equilibrium, departure from linkage disequilibrium, or null alleles in this population (Table S1), indicating that our chosen nine microsatellite markers were appropriate for matriline and patriline detection. The effective paternity number (*k*_e3_) was 3.23±0.32 SE using the equation of Nielsen et al. (2003).

There was no significant relationship between worker emergence date and patriline in any of the three colonies in experiment 2 (ANOVA: TK, n = 197, *F* = 0.690, *P* = 0.407; TG, n = 155, *F* = 0.005, *P* = 0.945; KM, n = 215, *F* = 3.023, *P* = 0.083). The survival time of the control was significantly longer than that of the workers infected with the two chosen fungal strains in all three colonies (log-rank test, *P* < 0.001). AIC values calculated in the full model using the results of experiment 2 (dependent variables: survival time and survival rate; predictor variable: each patriline, each strain, each emerged date and their interaction) in TK, TG, and KM colonies were 956.4, 756.1, and 1044.8. The result after performing stepwise model selection by AIC are presented in Table 2. We detected significant effects of patriline differences, emergence date, and the interaction of both on lifespan of *V. shidai* workers in the TK colony (Table 2). Similarly, we detected significant effects of patriline differences and strain differences on lifespan in the TG colony (Table 2). We detected significant effects of patriline differences and emergence date on lifespan in the KM colony (Table 2). The results of experiment 2 are illustrated in Fig. 2.

**Figure 2.**
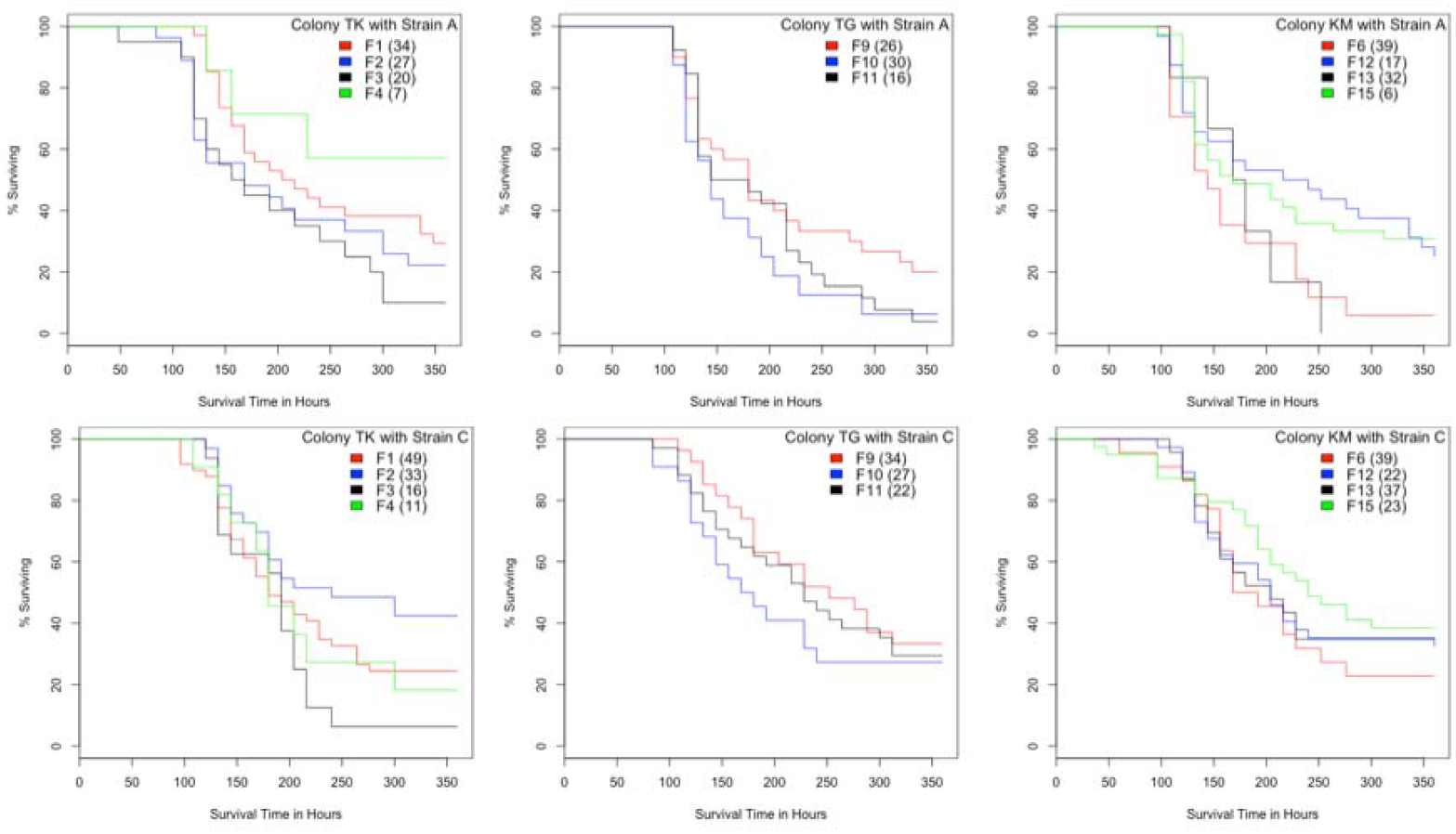
The survival curves of each patriline of each colony on the strain A and C in experiment 2. Legends show patrilines. Parentheses are number of samples. We showed results of statically analysis in Table 2.

**Table 2.** Cox survival analysis investigating the relationship between the lifespan and the strain, the patrilines, the emergence date, and those interactions.

| Dependent variable | Matriline | Predictor | Coef | Hazard ratio | SE | z | P |  |
| --- | --- | --- | --- | --- | --- | --- | --- | --- |
| **Lifespan** | TK, n = 197,<br>AIC = 956.4 | F3 vs F1 | -1.062 | 0.346 | 0.498 | -2.133 | 0.033 | * |
|  |  | F3 vs F2 | 0.212 | 1.236 | 0.559 | 0.379 | 0.704 |  |
|  |  | F3 vs F4 | -2.402 | 0.09 | 0.99 | -2.427 | 0.015 | * |
|  |  | strain | 1.171 | 3.227 | 0.662 | 1.771 | 0.077 |  |
|  |  | emergence date | 0.002 | 1.002 | 0.009 | 0.03 | 0.486 |  |
|  |  | F3 × emergence date vs F1 × emergence date | 0.105 | 1.111 | 0.081 | 1.295 | 0.195 |  |
|  |  | F3 × emergence date vs F2 × emergence date | -0.091 | 0.913 | 0.090 | -1.009 | 0.313 |  |
|  |  | F3 × emergence date vs F4 × emergence date | 0.277 | 1.319 | 0.137 | 2.019 | 0.043 | * |
|  |  | strain × emergence date | -0.152 | 0.859 | 0.093 | -1.630 | 0.103 |  |
|  | TG, n = 155,<br>AIC = 756.1 | F10 vs F9 | 0.274 | 1.315 | 0.221 | 1.237 | 0.216 |  |
|  |  | F10 vs F11 | 0.508 | 1.662 | 0.252 | 2.018 | 0.044 | * |
|  |  | strain | -0.608 | 0.544 | 0.193 | -3.150 | 0.002 | ** |
|  | KM, n = 215,<br>AIC = 1044.8 | F6 vs F12 | 0.629 | 1.876 | 0.237 | 2.650 | 0.008 | ** |
|  |  | F6 vs F13 | 0.153 | 1.166 | 0.207 | 0.740 | 0.359 |  |
|  |  | F6 vs F15 | 0.351 | 1.420 | 0.275 | 1.274 | 0.202 |  |
|  |  | emergence date | -0.084 | 0.919 | 0.030 | -2.828 | 0.005 | ** |

significance level \*, $P = 0.05$ , \*\*, $P = 0.01$

## Discussion

The five strains of *B. bassiana* used in this study all showed virulence to *V. shidai*,and for two of these the survival time of infected workers was significantly shorter than the control (Table S2). We also confirmed that *V. shidai* is polyandrous, with 3–4 major patrilines per colony, similar to other *Vespula* species (Goodisman et al. 2007; Bonckaert et al. 2008; Loope et al. 2014). Lifespan varied significantly both between colonies (= matrilines) and strains of *B. bassiana* (Table 1 and 2). Since *V. shidai* is an obligatory monogynous species, the observed differences in virulence between colonies suggests that matriline differences and/or the interaction of matriline and strain likely play a crucial role in colony resistance to parasites such as *B. bassiana*.

We also demonstrated that the lifespan of the workers infected with the fungus varied among patrilines, showing for the first time in eusocial wasps that resistance to pathogens varies depending on patriline. Such patriline differences have been shown in some bees and ants (Palmer and Oldroyd 2003; Baer and Schmid-Hempel 2003; Hughes and Boomsma 2004), as predicted by PPH; worker matrilines and/or patrilines probably have different levels of resistance to a single pathogen genotype. Pathogen variants show differences in virulence to individual matrilines or patrilines. Such variation in resistance among matrilines and patrilines would prevent colony failure as a whole due to reduced disease transmission among nestmates under genotype–genotype interaction in the host–parasite relationship (Sherman et al. 1988). There have been few reports that host survivorship depends on pathogen genotype (but see Barribeau et al. 2014). It is considered that the different strains of *B. bassiana* used in this study possessed different genotypes since they were isolated from different sources, and we demonstrated that the different strains varied in virulence, at least for some patrilines and matrilines. On this basis our results clearly support the PPH.

Dobelmann et al. (2017) compared natural viral infection rates and immune responses among patrilines within colonies of the polyandrous *V. vulgaris* but they found no differences due to patriline, perhaps due to lower sample sizes than those employed here. Our results suggest that the PPH likely explains why high colony genetic diversity is associated with high reproductive success in eusocial wasps, as found by Goodisman et al. (2007) and Dobelmann et al. (2017). As observed in other *Vespula* species (Ross 1986; Goodisman et al. 2007) we found that sperm mixing must have occurred in queens’ spermatheca because patrilines were evenly distributed across emergence dates resulting in increased genetic diversity within colonies.

In experiment 2, the environment in which the adult wasps were reared after emergence was under our control, but we did not control the environment of larval and pupal workers before emergence. In the KM colony, survival time of infected workers was dependent on emergence day, suggesting that resistance to pathogens is also affected by the environmental factors pertaining during the larval stage. However, even if some variation in survival is attributable to variation in the larval environment, this is almost certainly independent of the observed association with patriline.

Understanding the evolutionary processes that maintain polyandry in social insects has been one of the central challenges of social insect evolutionary biology in the past three decades (Mattila and Seeley 2007). Here for the first time we have demonstrated that the PPH is one of the evolutionary factors contributing to polyandry in eusocial wasps. Colonies we tested were shown to contain individuals of some patrilines that did not die after infection with *B. bassiana*. Patrilines conferring increased resistance may help a colony to maintain a minimum number of individuals to ensure colony survival and reproduction, and this may be particularly important in the early stages of colony development. *V. shidai* queens are known to usurp young nests of the same or other species (Saga et al. 2017). This intra- and interspecific usurpation by *V. shidai* leads to an increase in genetic diversity of nestmates in the early stages, and is likely to contribute to an increase in disease resistance. It is thought likely that increasing resistance to disease was a force that drove the evolution and maintenance of polyandry in eusocial insects.

We suggest that our methods should be applied to other species of wasp, since few have been studied. One might expect to find similar results in other obligately polyandrous Vespidae species, although the selective forces that favor facultative polyandry may be quite different (Loope 2015; Loope et al. 2017). Several other hypotheses (e.g., sperm limitation, Kraus et al. 2004; enhanced division of labor, Waibel et al. 2006; conflict reduction, Mattila et al. 2012, Loope 2015) have been proposed to explain the evolution of polyandry, and while they are not mutually exclusive with the PPH, all deserve further research.

## Acknowledgments

The authors would like to thank Katsuyuki Takahashi, Tsutomu Kamata and Kazuo Taguchi for helping me to collect and keep the wasp colonies. We sincerely thanks to Yoshikuni Hodoki for helping DNA analysis. We are very grateful to two anonymous reviewers for their valuable comments on previous versions of the manuscript. This study was supported in part by Takeda Science Foundation, Fujiwara Natural History Foundation, Funding of the Nagano Society for The Promotion of Science, Shimonaka Memories Foundation, Takara Harmonist Fund, Environmental research grant from Nissei Zaidan, and the Dream Project by Come on UP, Ltd.

## Notes

#### Summary of Updates

The title and main text have been minor changed.

