## Supplementary Material for "Polyandry and paternity affect disease resistance in eusocial wasps: support for the parasite–pathogen hypothesis"

### Genetic characteristics of the sample population

We confirmed that the microsatellite markers used in this study were suitable for paternity analysis. An inheritance data set (n = 10) was constructed in addition to the data set for the three colonies used for the pathogen application experiment; one individual was selected randomly from each of seven colonies of the same populations. These data sets were tested for deviation of Hardy–Weinberg equilibrium and linkage disequilibrium using Genepop 4.4(Rousset 2008). In addition, these data sets were also tested for null allele using the software Cervus 3.0.

The genetic characteristics of the microsatellites in this population are summarized in Table S2. No linkage disequilibrium was observed in this population (*p* > 0.05), no significant loss of Hardy–Weinberg equilibrium was observed (*p* > 0.05) and no null allele was detected (Sequential Bonferroni, *p* > 0.05). On this basis, the nine microsatellite loci used in this study were considered to be appropriate for paternity analysis.

| Table S1. Genetic characteristics of the nine microsatellite markers in the Nakatsugawa population | | | | | |
| --- | --- | --- | --- | --- | --- |
| Locus | *n* | *Na* | *Ho* | *He* | *F (Null)* |
| Rufa5  Rufa19  List2001  List2003  List2004  List2019  List2020  VMA3  VMA6 | 10  10  10  10  10  10  10  10  10 | 3  7  6  9  9  5  4  10  5 | 0.900  0.900  0.800  1.000  1.000  0.700  0.700  0.900  0.900 | 0.679  0.826  0.800  0.889  0.874  0.626  0.695  0.889  0.789 | −0.166  −0.068  −0.026  −0.089  −0.099  −0.093  −0.044  −0.029  −0.090 |
| n, number of samples; Na, number of alleles; *Ho*, observed heterozygosity; *He*, expected heterozygosity; *F (Null)*, results of test for null alleles | | | | | |

| **Table S2**. Survival time (in days) and survival rate of workers at 7 days after each treatment. | | | | | |
| --- | --- | --- | --- | --- | --- |
| Colony no. | Strain of fungus | Sample size | Survival time (days) | Survival rate^$^ | *P* value |
| 1 | Control | 10 | 5.1 ± 2.2 | 0.50 |  |
|  | A | 20 | 3.5 ± 1.5 | 0.00 | 0.018^#^ |
|  | B | 20 | 4.1 ± 1.1 | 0.00 | 0.062 |
|  | C | 20 | 4.1 ± 1.9 | 0.00 | 0.022^#^ |
|  | D | 22 | 4.7 ± 0.6 | 0.00 | 0.231 |
|  | E | 22 | 4.5 ± 1.4 | 0.00 | 0.062 |
| 2 | Control | 10 | 4.5 ± 2.0 | 0.20 |  |
|  | A | 14 | 3.2 ± 1.2 | 0.00 | 0.011^#^ |
|  | B | 14 | 3.0 ± 1.5 | 0.00 | 0.015^#^ |
|  | C | 14 | 4.1 ± 1.6 | 0.00 | 0.346 |
|  | D | 18 | 3.3 ± 1.6 | 0.00 | 0.047 |
|  | E | 20 | 3.6 ± 1.5 | 0.00 | 0.075 |
| 3 | Control | 10 | 4.5 ± 1.8 | 0.20 |  |
|  | A | 20 | 2.9 ± 1.1 | 0.00 | 0.009* |
|  | B | 20 | 2.3 ± 1.1 | 0.00 | 0.0015* |
|  | C | 20 | 3.4 ± 1.2 | 0.00 | 0.058^#^ |
|  | D | 23 | 2.4 ± 1.3 | 0.00 | 0.0033* |
|  | E | 21 | 3.2 ± 1.4 | 0.00 | 0.032^#^ |
| 4 | Control | 10 | 5.2 ± 1.3 | 0.30 |  |
|  | A | 18 | 2.9 ± 1.1 | 0.00 | 0.0028* |
|  | B | 16 | 3.7 ± 1.4 | 0.00 | 0.030^#^ |
|  | C | 18 | 3.7 ± 1.6 | 0.00 | 0.037* |
|  | D | 18 | 3.4 ± 1.2 | 0.00 | 0.0011* |
|  | E | 18 | 3.4 ± 1.4 | 0.00 | 0.0076* |
| 5 | Control | 10 | 2.9 ± 1.9 | 0.10 |  |
|  | A | 11 | 3.0 ± 1.3 | 0.00 | 0.958 |
|  | B | 11 | 3.6 ± 1.4 | 0.00 | 0.350 |
|  | C | 10 | 3.5 ± 2.1 | 0.00 | 0.579 |
|  | D | 13 | 3.8 ± 1.5 | 0.00 | 0.312 |
|  | E | 9 | 3.3 ± 1.7 | 0.00 | 0.794 |

Values are mean ± SD. Survival curves between control and the workers infected with each strain (A–E) were compared using log-rank test and the calculated *P* values are shown in the table. ^#^, marginally significant (α = 0.1); *, significant (α = 0.05) after sequential Bonferroni's correction; ^$^, survival rate at the 7th day after treatment.
